# ΦX174 Attenuation by Whole Genome Codon Deoptimization

**DOI:** 10.1101/2020.02.10.942847

**Authors:** James T. Van Leuven, Martina M. Ederer, Katelyn Burleigh, LuAnn Scott, Randall A. Hughes, Vlad Codrea, Andrew D. Ellington, Holly Wichman, Craig Miller

**Affiliations:** Department of Biological Science, University of Idaho, Moscow, ID 83844; Institute for Modeling Collaboration and Innovation, University of Idaho, Moscow, ID 83844; Seattle Children’s Research Institute, Seattle, WA; Applied Research Laboratories, University of Texas, Austin, TX 78758; Biotechnology Branch, CCDC US Army Research Laboratory, 2800 Powder Mill Rd, Adelphi, MD 20783; Institute for Cellular and Molecular Biology, University of Texas, Austin, TX 78712

## Abstract

Natural selection acting on synonymous mutations in protein-coding genes influences genome composition and evolution. In viruses, introducing synonymous mutations in genes encoding structural proteins can drastically reduce viral growth, providing a means to generate potent, live attenuated vaccine candidates. However, an improved understanding of what compositional features are under selection and how combinations of synonymous mutations affect viral growth is needed to predictably attenuate viruses and make them resistant to reversion. We systematically recoded all non-overlapping genes of the bacteriophage ΦX174 with codons rarely used in its *E. coli* host. The fitness of recombinant viruses decreases as additional deoptimizing mutations are made to the genome, although not always linearly, and not consistently across genes. Combining deoptimizing mutations may reduce viral fitness more or less than expected from the effect size of the constituent mutations and we point out difficulties in untangling correlated compositional features. We test our model by optimizing the same genes and find that the relationship between codon usage and fitness does not hold for optimization, suggesting that wild-type ΦX174 is at a fitness optimum. This work highlights the need to better understand how selection acts on patterns of synonymous codon usage across the genome and provides a convenient system to investigate the genetic determinants of virulence.

## Introduction

### Synonymous compositional features of viral genomes

The unequal use of synonymous codons is known as codon usage bias (1). Codon biases are the result of an interaction between mutational and selective pressures (2, 3). Understanding how these two forces combine to determine codon usage in a genome is challenging. A commonly accepted explanation is that the rate of translation (i.e. the amount of proteins being made) is affected by the abundance of tRNAs that pair with codons on a strand of mRNA. Codons that result in the optimal amount of protein then have a selective advantage. However, there are many other compositional features within the DNA sequences of protein coding genes upon which selection acts. These features include: the genic GC content (4–6), CpG or TpA dinucleotides (7–11), codon pairs (12–15), endonuclease recognition sites (16–19), intron splicing motifs (20), mRNA folding stability (21–25), ribosomal pausing sites (26, 27), concentration of non-preferred codons at transcripts 5’ end (28–30), autocorrelation of codons on transcript (31), and capacity of codon order to influence co-translational folding of proteins (32, 33). At any given site, natural selection may act on one or more of these features to favor the use of certain codons over other synonymous ones (34). While the strength of selection on synonymous mutations is generally considered weak to neutral and genetic drift should therefore also exert a strong influence, the overall signature of selection in the form of non-random codon usage is broadly observed across life (34–36) where selection has been shown to strongly favor one synonymous mutation over others (37–40).

As viruses must utilize their hosts’ cellular machinery, there is an expectation that virus genomes are enriched for host-preferred codons. This appears to be only partially true. Many viral genomes contain more host-preferred codons than expected by chance, especially in highly expressed genes encoding viral structural proteins (41). However, many viral genes are not enriched in host-preferred codons. Sometimes unpreferred codons are used to regulate viral gene expression (42). Other virus genomes appear to have little preference for codons abundant in the host genome. For instance, Lucks et al. found that the majority of 74 bacteriophage genomes show no significant preference for host-preferred codons (41). Similar discordance between host and viral codon usage patterns are observed in other studies (43–45). This discordance could be caused by insufficient selection on codon usage, host-phage relationships that are too short-lived for selection to fine tune codon usage in the phage, or inadequate understanding of the features under selection.

### Vaccine development by synonymous recoding

Empirically developed (e.g. serial passage viral adaptation) vaccines have saved millions of lives over the last century, yet methodological improvements make rationally designed, recombinant vaccines attractive because they can be rapidly produced and specifically engineered for safety and effectiveness (46–50). One proposed method of generating recombinant vaccines involves making many synonymous, attenuating changes to viral genomes, i.e. “deoptimizing” the viral genes (51). The recombinant vaccine can be made by either editing the genome of the wild-type virus or by generating a viral genome entirely from synthesized nucleic acids. Synonymous deoptimization offers a potentially efficient and effective way of making vaccines: the protein sequences of recoded vaccines are identical to their target viruses, they replicate in their host to provide prolonged exposure to the antigen, and the introduction of many synonymous changes presumably assures evolutionary robustness, preventing the evolution of virulence by reversion.

Poliovirus serves as a very good example for the synonymous recoding strategy. Development of a robust, live-attenuated poliovirus vaccine is desired because in some areas on Earth wild poliovirus and the emergent vaccine-derived polioviruses (cVDPVs) continues to cause concern over the resurgence of poliomyelitis (52–55). A synthetic poliovirus was assembled in 2002 (56), codon deoptimized in 2006 (57, 58), codon pair deoptimized in 2008 (59), and dinucleotide deoptimized in 2009 (11). In all cases, attenuated viruses were produced by recoding the P1/capsid region of the genome. *In vitro*, these viruses replicate slower and produce lower viral titers than wild-type virus. *In vivo* the codon pair deoptimized strain protected mice against challenge by wild poliovirus (59). While the mechanism of attenuation is not yet fully elucidated, reduced protein expression of the deoptimized genes is observed (11, 57–59). These deoptimized poliovirus constructs are genetically stable and remain non-virulent for up to 25 passages in cell culture (57, 59).

The apparent success of building poliovirus vaccine candidates using synonymous recoding led to similar attempts to develop vaccines for influenza, adeno-associated, human immunodeficiency, papilloma, chikungunya, respiratory syncytial, simian immunodeficiency, porcine reproductive and respiratory syndrome, echovirus 7, tick-borne encephalitis, vesicular stomatitis, dengue, T7, Lassa, adeno, and swine fever viruses (reviewed in (60)). The most common method for synonymously deoptimizing viruses is recoding wildtype genes with increased proportions of non preferred codons (58, 61–65, 65–69) although other methods of recoding have been successful as well. For example, viral fitness was decreased when synonymous substitutions were randomly introduced (70–72), when codons were replaced by those infrequently used in viral (not host) genes (57, 66, 73), when the proportion of optimal codons was *increased* (64, 74, 75), or when the number of codons one substitution away from a translational termination codon was increased (76).

### No predictive understanding of synonymous recoding

While it is clear that synonymous recoding causes attenuation and the strategy holds promise for vaccine development, we lack a predictive understanding of the process. Part of this results from the biological complexity and variation in the systems involved. In many cases, the fitness impact of recoding is cell-line dependent (62, 66, 68–70, 77–79), is inconsistent between *in vivo* and *in vitro* experiments (63, 69, 79, 79), or is temporally variable (73). Another obstacle is the nature of the genetic code itself: it is generally challenging to manipulate one compositional feature of the genome and hold all the others fixed. For example, when codons are shuffled to change codon pair frequency, mRNA stability may be affected. Or when codons are deoptimized, codon pair frequencies also change. This makes it difficult to attribute the cause of fitness decreases to one factor (*e.g.* codon usage adaptation), especially when the features are correlated. As we do here, most studies have focused on manipulating a single compositional feature of the genome and measuring its impact on fitness. Standardizing recoding methodologies and features measured across studies would greatly improve our understanding of the factors that drive fitness decreases and other phenotypic effects caused by synonymous deoptimization.

Despite the optimistic results achieved in studies on synonymous recoding to date, basic questions underlying the method itself remain unanswered (60). What is the best strategy to perform synonymous recoding to achieve attenuation? Can generalities be made about the extent of recoding and the degree of attenuation—or will the biological details and idiosyncrasies of each system preclude this? What is the mechanistic cause of attenuation from synonymous recoding? Are viruses recoded this way robust against fitness recovery? Is it more effective to maximally recode less of the genome (say one gene), or make the recoding less severe, but distribute it across the genome? As an increasing proportion of the genome is recoded, or equivalently, as multiple recoded parts are combined, does attenuation respond in an additive or non-additive manner? A deeper understanding of genome evolution and synonymous sequence choice is required to answer these questions.

In this paper, we focus on two issues related to the codon deoptimization of viruses. First, we seek to compare the fitness effects of mutating different genes in the same virus. Second, we seek to understand how, when attenuating mutations are combined, they interact to affect fitness (*i.e.* epistasis of deleterious mutations). The nature of epistasis among fragments is crucial for modeling fitness effects: if mutational effects combine synergistically (*i.e.* the combined fitness being even lower than predicted from the observed individual effects), the range in the number of mutations needed to achieve the targeted attenuation level would tend to be reduced (fig. 1). Conversely, if they combine antagonistically (*i.e.* the combination of mutations are less attenuated than predicted from individual effects), it may be easier to achieve a target attenuation level, but there may be a limit to how much attenuation is possible. If mutations, in combination, display sign epistasis, irregular magnitude epistasis, or even vary between synergistic and antagonistic epistasis, it will suggest the underlying process is complex and difficult to predict and generalize. To evaluate these issues, herein we have recoded all the non-overlapping genes of the bacteriophage ΦX174 in fragments and combined recoded fragments in all possible within-gene permutations, and measure the fitness of the resulting recoded bacteriophage.

**Fig. 1.**
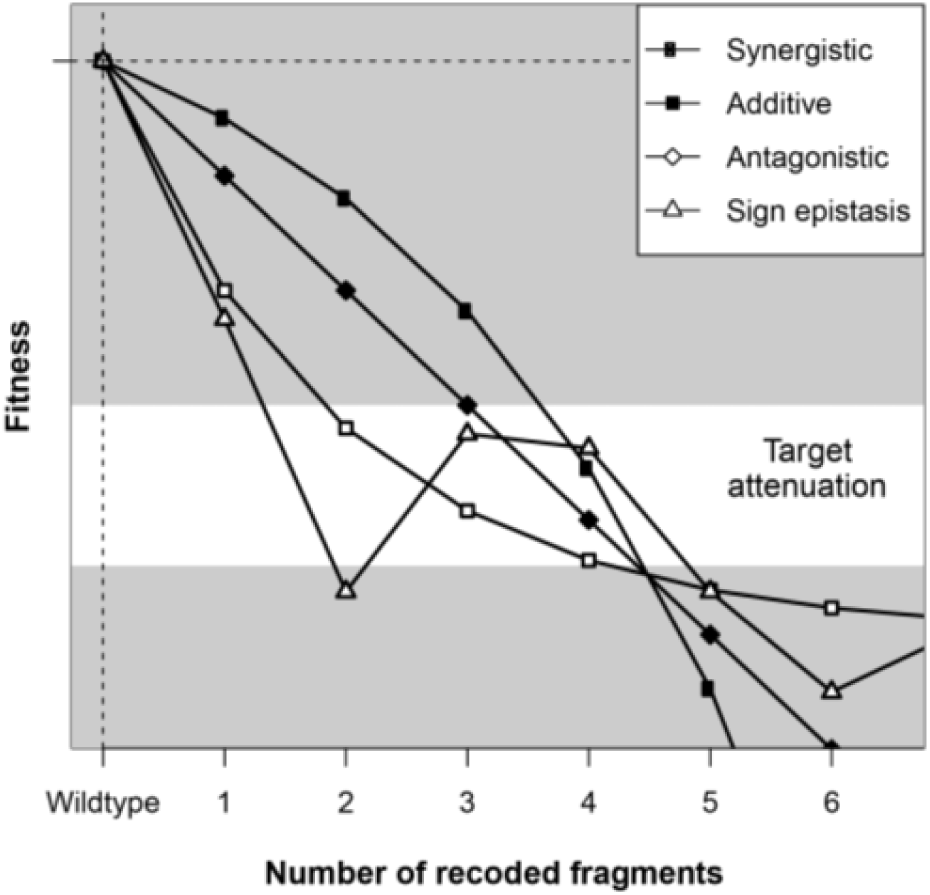
Hypothetical data illustrates that epistasis determines the number of mutations needed to attenuate a virus. The “targeted attenuation” window is more difficult to achieve if mutational effects combine synergistically than if mutations combine antagonistically because the slope of the curve is steeper (fewer mutations in the window). Notice that this pattern is reversed if a slight level of attenuation is desired (near y=0). If sign epistasis, or even irregular magnitude epistasis is observed, then the underlying nature of interactions are more difficult to predict and generalize.

## Results

### Synonymously deoptimizing ΦX174 genes

ΦX174 is a bacteriophage with a 5.4kb single-stranded DNA genome containing 11 genes (fig. 2a, table 1). We measured codon usage bias of ΦX174 genes using the codon adaptation index (CAI). CAI is a gene-level statistic running from zero to one that summarizes the extent that codons in a gene are used, rarely (CAI nearer 0) or commonly (CAI nearer 1) among highly expressed host genes (80). We found that most ΦX174 genes are not particularly enriched for preferred *E. coli* codons (fig. 2b). Only gene J has a codon adaptation index (CAI) value equivalent to highly expressed *E. coli* genes. Gene K uses the most unpreferred codons and has a CAI value near the lowest value observed for genes in the *E. coli* genome. All other ΦX174 genes have CAI values within the range of most *E. coli* protein-coding genes. ΦX174 structural proteins (B, D, F, G, H, J) have higher CAI values than non-structural genes (A, C, E, K), suggesting that high expression of these proteins is important for viral fitness. When we computationally deoptimized entire ΦX174 genes (*i.e.* recoded them to use the least-preferred codons throughout), the resulting CAI values were in the lower tail or even below the tail for all *E. coli* genes (fig. 2, supplementary table S1). These reductions in CAI were the result of changing between 42% (20/48 codons for gene C) and 75% (24/32 codons for gene J) of the codons of a gene, corresponding 15-32% of its base pairs (supplementary table S2). All the other ΦX174 genes fall within this range of recoding. We calculated additional metrics of codon adaptation including an alternative version of CAI (81), tRNA adaptation index: tAI (82), index of translation elongation: I_TE_ (83), relative codon adaptation: RCA (84), the number of effective codons: Nc (85), and the starvation codon adaptation index: sCAI (86). Nc differs from the other indices in that it measures the deviation from uniform codon usage and does not score codon preference. The other indices compare the codon usage of any given gene to a set of genes that are known to be highly expressed in a cell. These latter indices differ from one another in how codon optimality is scored (*e.g.* taking into account tRNA gene abundances). Calculation details are in the Materials and Methods. All analyses produced qualitatively similar rank-orders for ΦX174 genes (supplementary table S1).

**Table 1.** ΦX174 gene function and protein copy number required for the assembly of one virion.

| Gene | Protein Function | Copies | Gene length (bp) |
| --- | --- | --- | --- |
| A | replication initiation, viral strand synthesis | 30 | 1,541 |
| A* | competitive exclusion | 30 | 1,025 |
| **B** | **capsid morphogenesis** | 60 | 362 |
| K | unknown, not essential |  | 170 |
| C | DNA packaging | 60 | 260 |
| **D** | **capsid morphogenesis** | 240 | 458 |
| E | cell lysis |  | 275 |
| **J** | **core protein, DNA condensation** | 60 | 116 |
| **F** | **major capsid protein** | 60 | 1,283 |
| **G** | **major spike protein** | 60 | 527 |
| **H** | **minor spike, pilot protein** | 12 | 986 |

<sup>a</sup> Genes encoding structural proteins are bolded.

**Fig. 2.**
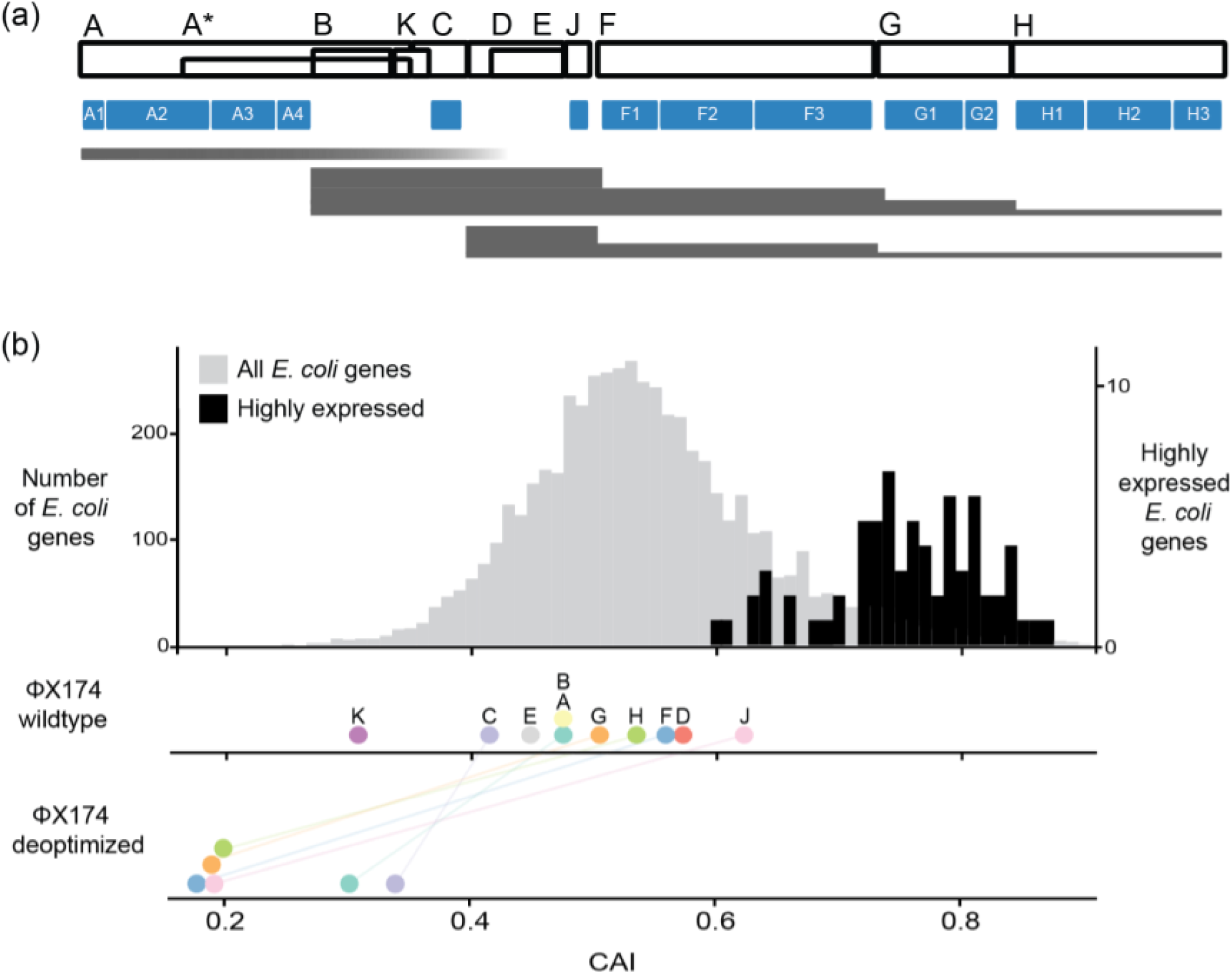
ΦX174 genome organization and capacity for deoptimization relative to host genes. (a) Genes on the ΦX174 genome are shown as empty black boxes. Recoded regions are shown as filled blue boxes. These fragments are named consecutively (e.g., A1, A2, A3, A4, F1, F2, F3,…). Approximate transcript expression levels are shown as filled grey bands whose heights are proportional to the relative number of transcripts by RT-qPCR. (b) Codon adaptation index (CAI) of *E. coli* genes, genes in wild-type ΦX174, and deoptimized ΦX174 genes. Genes highly expressed in *E. coli* are rescaled to the secondary y-axis.

### Codon deoptimization of ΦX174 genes reduces viral fitness

We codon deoptimized the genes of ΦX174 by recoding nearly all synonymous residues to the least preferred *E. coli* codons. We did not recode regions of genes that overlapped with other genes nor the first six residues of each gene since these residues are known to have strong effects on gene expression (87). Four of the six recoded variants were less fit than wild type ΦX174 (fig. 3a). The construct containing the fully deoptimized G gene could not be recovered, even after growing the strain overnight on the susceptible host cells in an attempt to obtain a recovery mutation. Recoding highly expressed genes (J, F, and G) resulted in larger fitness decreases than recoding lowly expressed genes (A and H). Although the number of variants built was small, the fitness effects of deoptimization were correlated to the proportion of codons edited and the CAI of the recoded genes (fig. 3b and supplementary table S2).

**Fig. 3.**
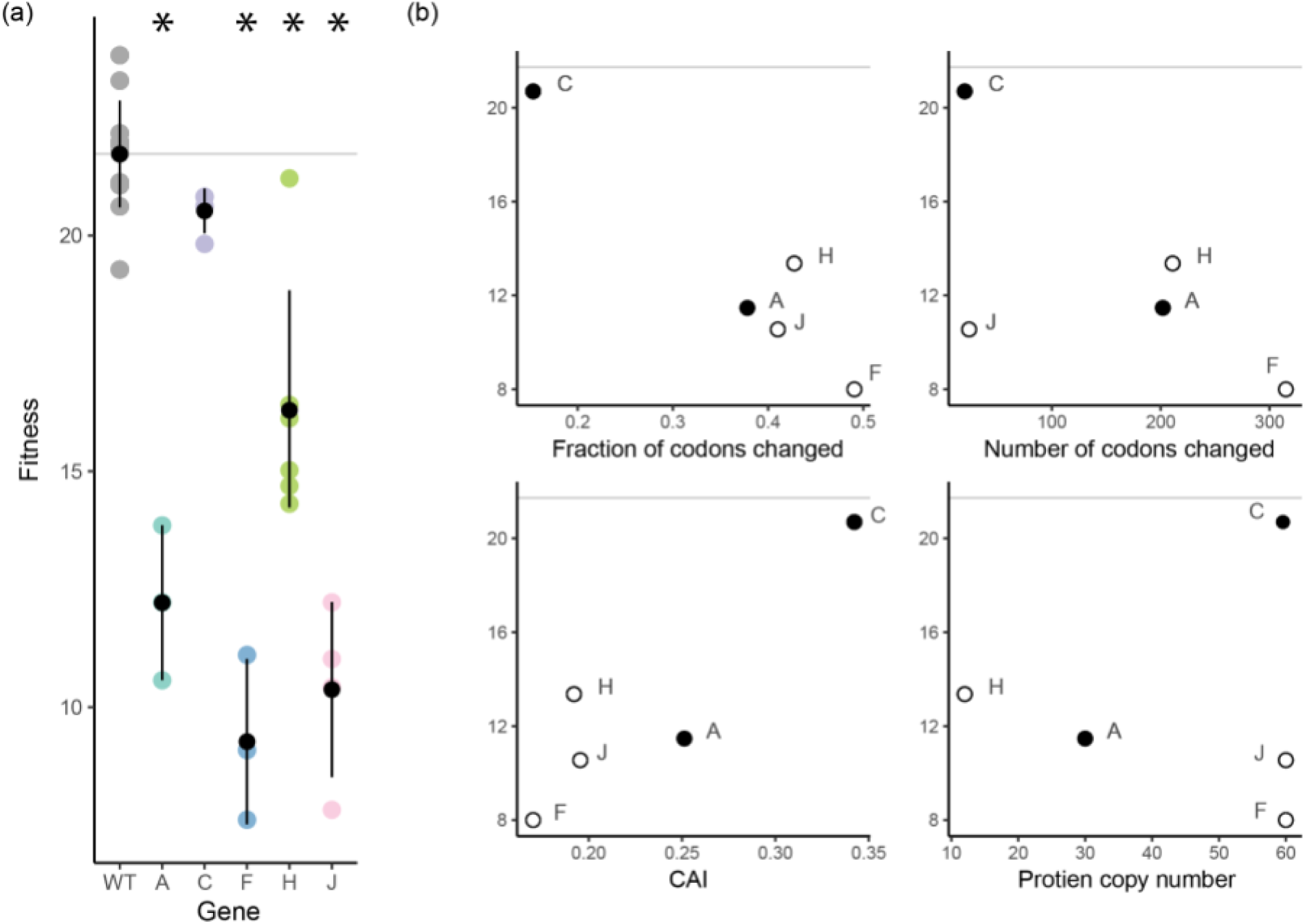
Fitness effects of deoptimizing ΦX174 genes. (a) The fitness of wild type and deoptimized ΦX174 strains containing recoded genes are shown in replicate. Means and standard error bars are shown in black. Fitness values that are significantly different from wild type are indicated with asterisks. (b) Fitness is plotted against measures that potentially explain fitness decreases. Protein copy number is the number of proteins needed to virion assembly (see Table 1). Fitness is the number of doublings per hour (log2 of the ratio of the phage concentration at 60 min divided by the phage concentration at time zero). The fitness of wild-type is indicated with a grey horizontal line. Structural proteins are shown as empty circles. Deoptimizing gene G yielded no viable phage.

### Reconstructing a combinatorial fitness landscape for deoptimized genes

We segmented ΦX174 genes into gene fragments (fig. 2) and measured the fitness of all of the possible within-gene combinations of deoptimized fragments (fig. 4 and supplementary table S3). Of the 12 deoptimized strains with only one deoptimized fragment, only 6 have fitness values below wild type. The moderate fitness effects of these partially recoded genes allowed us to observe how deleterious effects combine as compared to the more deleterious effects observed when deoptimizing complete genes. As additional deoptimized fragments are joined, the fitness of the resulting viruses decrease (fig. 4b). In most cases combining deoptimized fragments results in less fit viruses. The exception is gene A where instances of sign epistasis are observed. Specifically, the average fitness of A1+A3, A2+A3, A1+A3+A4, A1+A2+A4, A2+A3+A4, and A1+A2+A3+A4 are all higher than at least one of their constituent fitness values (fig. 4b). To further investigate how deleterious effects combine, we employed a statistical framework for calculating the best-fitting model of epistasis.

**Fig. 4.**
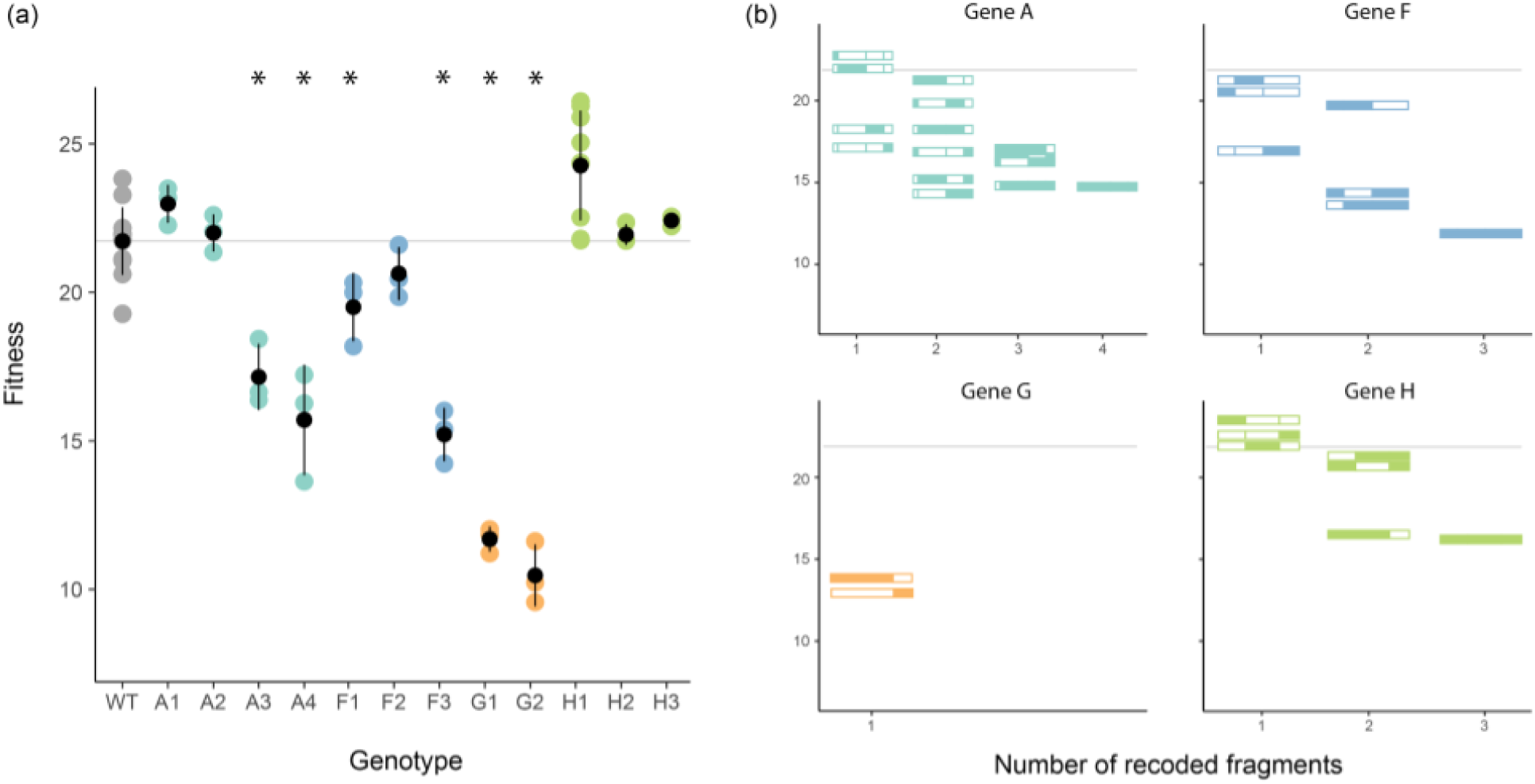
Fitness of ΦX174 when deoptimized gene fragments are combinatorially joined. The fitness of variants containing one deoptimized fragment (a) and all possible within-gene variants (b) was measured and compared to wild type (grey horizontal line). In (b), fragment lengths are drawn to scale. Filled colors indicate the deoptimized fragments while unfilled blocks indicate wild-type fragments. In both (a) and (b), fitness is shown as log2 fold increase in the number of phage per hour.

### Fitting models of epistasis to combinatorial fitness data

The combinatorial network of genotypes that we generated in this work can be analyzed by applying simple models of epistasis (88) to determine how the effects of mutations combine. We fit the data for genes A, F and H to three basic models—additive, multiplicative, and stickbreaking—which gave rise to no, antagonistic, and synergistic epistasis respectively (see figure 1 and (89)). In fitting the three models, we conducted two analyses for each gene: one of absolute fit where we assess if the data is consistent with each model individually, and one of relative fit wherein one of the three models is assumed to be correct. The results from this analysis were not highly conclusive, but suggest the nature of epistasis is heterogeneous across different genes. For genes F and H, none of the three models could be rejected based on absolute goodness of fit (table 2). For gene F, the additive model provides the best fit to the data. For gene H, stickbreaking gives the best fit (R^2^=0.885), consistent with synergistic epistasis. This is visually clear in figure 4b, where the fully recoded gene H (3 recoded fragments) has far lower fitness than one would expect based on the individually recoded fragments—all of which were basically neutral.

**Table 2.** Models of epistasis fit to combinatorial fitness data.

| Gene | P-value (absolute fit)<sup>a</sup> |  |  | Posterior (relative fit)<sup>b</sup> |  |  | R<sup>2</sup> |  |  |
| --- | --- | --- | --- | --- | --- | --- | --- | --- | --- |
|  | Add | Mult | Stick | Add | Mult | Stick | Add | Mult | Stick |
| A | 0.011 | 0.066 | <0.001 | 0.145 | 0.845 | 0.009 | 0.761 | 0.860 | 0.388 |
| F | 0.388 | 0.071 | 0.235 | 0.609 | 0.090 | 0.301 | 0.965 | 0.843 | 0.950 |
| H | 0.513 | 0.489 | 0.733 | 0.056 | 0.054 | 0.891 | 0.709 | 0.562 | 0.885 |

Note - A regression of each recoded fragment's fitness effect (against background) under each model was performed. The p-values of each regression were combined by taking the sum of their logs. Using parametric bootstrap, the distribution of this sum was simulated. The overall p-value is estimated by the proportion of simulations where the sum of logs is ≤ the observed value.
<sup>a</sup> Absolute goodness of fit. Small p-values indicate that the data is inconsistent with the model (gray-filled). When a model is correct, a recoded block's effect is uncorrelated to background fitness. The p-value indicates how often, under parametric bootstrapping, the correlation of effect to background fitness is as strong as or stronger than that observed in the real data.
<sup>b</sup> The posterior probability assumes that one of the three models—additivity, multiplicative, stickbreaking—is correct.

Visually, a pattern of antagonistic epistasis was observed for gene A, as several of the variants with two recoded fragments had fitnesses as low or even slightly lower than the three- and four-recoded fragment variants (fig. 4b). Indeed, the additive and stickbreaking models were rejected for gene A based on absolute goodness of fit (table 2). The multiplicative model, with its antagonistic pattern of epistasis, was not rejected, but the p-value was marginal (p=0.066). Strong antagonistic epistasis is occuring for gene A—even stronger than that predicted under the multiplicative model. This was revealed by regressing background fitness against fitness effect (fig. 5a and supplementary figs. S1-S3). When effects were measured as differences (the additive model), negative/positive slopes corresponded to antagonistic/synergistic epistasis. Under the correct model, no correlation exists and slopes are expected to be random deviations around zero. Under the additive model (fig 5a), a clear pattern across all four recoded fragments where the effect of the fragment becomes more strongly deleterious on higher fitness backgrounds (negative regression slopes) was observed. When the p-values of the individual fragments were combined, their result is significant (supplementary fig. S1). The analogous regression under the multiplicative model was less extreme, but even here the slopes were consistently negative, indicating a level of antagonistic epistasis beyond multiplicative (supplementary fig. S2).

**Fig. 5.**
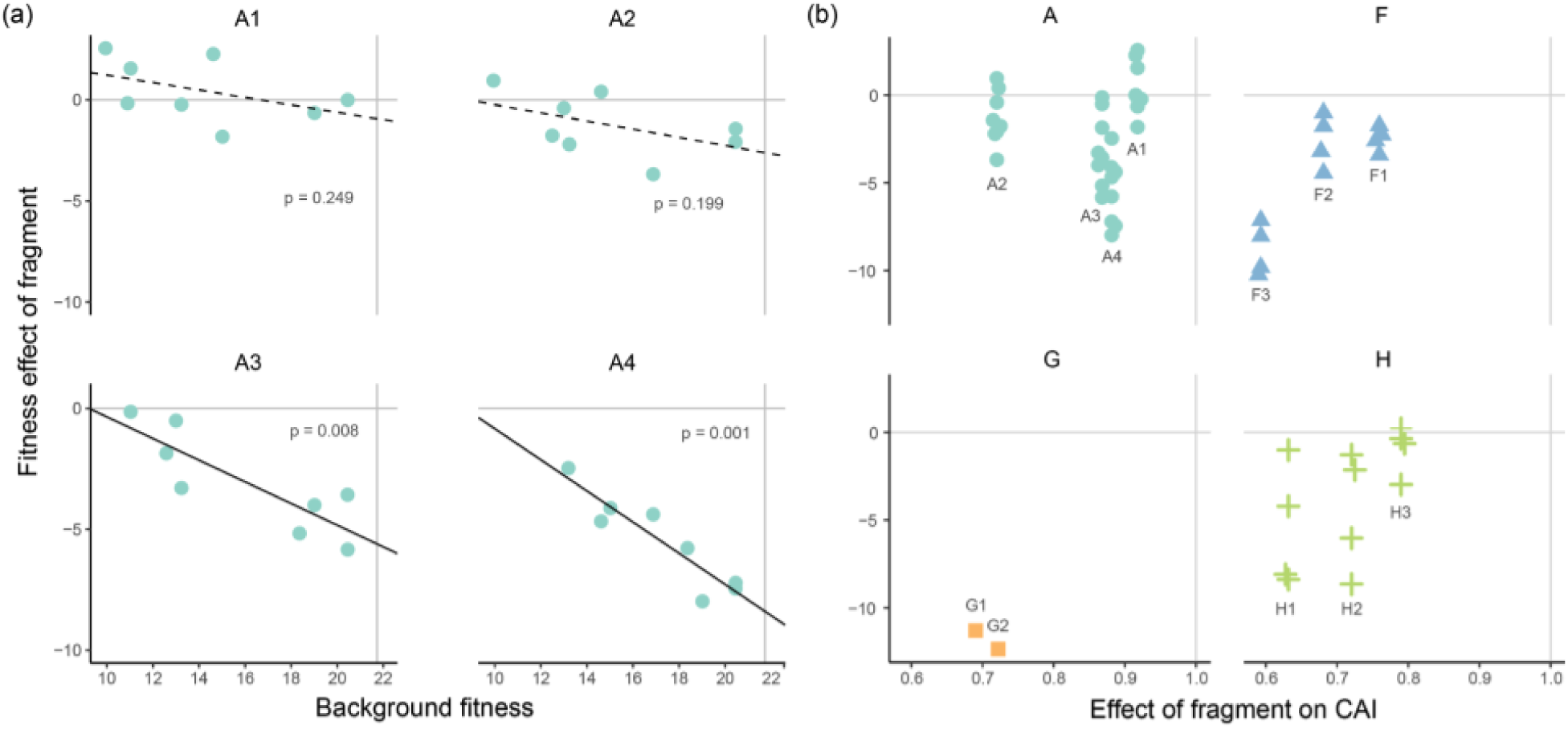
Fitness effect of each recoded fragment on all possible backgrounds. (a) Regressions of each recoded fragment’s fitness effect under the additive model against background fitness. The y=0 line is a perfect fit to the additive model with no residual effects of background. The p-value for each linear regression is shown. The overall fit of every model for each gene is shown in Table 3. In (b), the effect a fragment has on CAI is shown against its effect on fitness from the background to the background plus the recoded fragment (A2 -> A1+A2, A2+A3 -> A1 + A2+A3) is shown for all fragments. Slight point jitter was used for visualization.

### Correlating codon deoptimization to combinatorial fitness data

Ultimately, our goal was to correlate changes in genomic properties (*e.g.* codon preference) to changes in viral fitness. The most straightforward method of analysis would be to regress the two measurements, and indeed the fitness of deoptimized variants was linearly correlated to CAI (R^2^=0.36, p=2E-16, fig. 6a). However, it is worth noting that the data points used in this regression were not independent because the deoptimized fragments were combined to achieve higher levels of deoptimization. Our combinometic method of making variants also allowed us to correct for the cumulative fitness effect of combined fragments by calculating the effect of adding any particular fragment to different backgrounds (fig. 5b). For example, the effect of deoptimizing the F1 fragment was measured by comparing the fitness values of WT to deoptimized F1 (20-21 = −1), or F2 to F1+F2 (18-20 = −2), or F3 to F1+F3 (14-16 = −2), or F2+F3 to F1+F2+F3 (8-10 = −2). Thus, deoptimizing F1 resulted in an average fitness effect of about -2. This background subtraction approach corrected for the non-independence of data points in regressions between change in fitness from WT and change in CAI from WT.

**Fig. 6.**
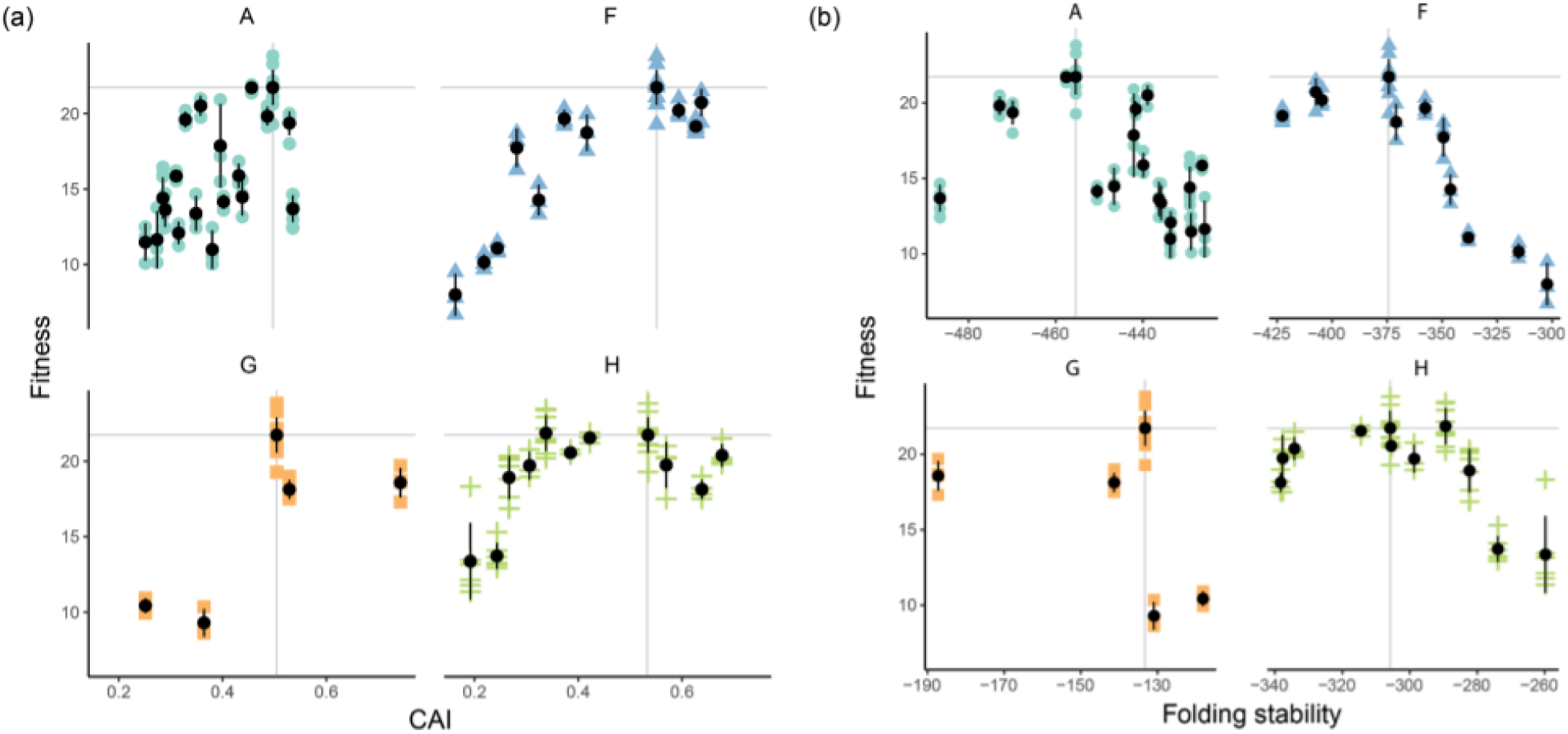
Fitness of recoded viruses correlates with codon usage bias and mRNA folding stability. (a) Codon usage bias (CAI) compared to fitness for viruses optimized and deoptimized in genes A, F, G, and H. Fitness and CAI of wild type are indicated with grey horizontal and vertical lines. Points to the right of these lines are optimized. Points to the left are deoptimized, (b) Viral fitness compared to mRNA folding stability (mfold). Wild type values are indicated with grey horizontal and vertical lines. Less stable transcripts (deoptimized genes) have less negative values and are right of wild type. More stable transcripts (optimized genes) have more negative values.

When we applied this correction, we observed a wide variance in fitness effects (fig. 5b). For example, in some backgrounds, adding deoptimized H1 reduced fitness by only ~1 doublings/hr. In other backgrounds, H1 reduced fitness by ~8 doublings per hour. Despite this variation, there is a good correlation between change in CAI and change in fitness (R^2^=0.58, p=4.8E-10, fig. 5 and supplementary table S4). Applying this background correction indicates that only a portion of the fitness changes can be explained by changes in codon usage bias. This is particularly true for genes A and H. Fragments in gene F seem to have more consistent effects (fig. 5b).

### How different synonymous features correlate with fitness

We replaced ΦX174 codons with less preferred codons without consideration for how alterations might affect other features in the genome. As mentioned in the introduction, many such features may be under selection. To investigate unintended consequences of codon deoptimization, we calculated numerous genome characteristics to see if any correlate with the fitness decreases observed in deoptimized fragments (supplementary table S1). We included many different measures of codon usage bias (CAI, tAI, ITE, etc), codon pair bias (CPB), frequency of Shine-Dalgarno motifs, mRNA folding stability, as well as simply the number of changes made. The best predictor of fitness is the folding stability of the codon deoptimized mRNA (R^2^=0.67, p.adj=0.02), which performed significantly better than the best measure of codon usage bias which was CAI using the Xia2007 method (ΔAIC=15.5). This correlation is easily observed when mRNA stability values are plotted against change in fitness (fig. 6b). We were interested to see if the correlation between genomic features like CAI and fitness held up even when features were optimized, so we replaced ΦX174 codons with codons frequently used in *E. coli* (supplementary table S2), expecting to observe beneficial fitness effects. However, in all cases, fitness was either unaffected or reduced (fig. 6a and supplementary table S3). Because of this, if these optimized constructs are included in the regression models, the number of sites changed and fraction of gene edited become the metrics that best predict fitness from genomic measures. We observed a peak-shaped fitness landscape when combining the optimized and deoptimized data set; this is discussed below.

## Discussion

### Patterns of synonymous codon usage biases

Synonymous codon usage biases are present in genomes across the tree of life (35). We often think of these biases as having little consequence during the natural evolution of organisms because the strength of selection acting on any one synonymous mutation is generally weak. Nevertheless, the presence of biases shows that selection acts with sufficient strength to maintain them in the face of genetic drift. The prevailing theories on the preservation of codon biases suggest that codon choice is primarily driven by selection on translational speed and mRNA stability (1, 15, 35, 90). The enrichment of codons that use abundant tRNAs in highly expressed genes points towards a model where translation speed is correlated to tRNA abundance. We find that the most highly expressed ΦX174 gene ranks second best in its use of host-preferred codons, but only marginally better than the average *E. coli* gene (fig. 2). That the majority of ΦX174 genes fall below average in preferred codon usage bias according to *E. coli* usage patterns is not surprising—many viruses do not favor the most preferred host codons. A myriad of reasons may explain why viral genes do not evolve to their full codon usage potential. Assuming a selective explanation, it could be that codon usage controls the stoichiometric ratio between viral genes (91), temporally regulates gene expression (42, 73, 92), facilitates co-translational folding (33), dampens protein expression to avoid host immune responses (64, 93), or is limited by other compositional features. Comparative genomics between ΦX174, G4, α3, and ΦMH2K suggest that the codon usage biases are a conserved feature of these microviruses (94). Our data cannot point towards any one of these explanations, although we note that the high-copy number proteins of ΦX174 tend to have higher CAI values and are more easily deoptimized.

### Recoding of ΦX174

Synonymous mutations that disrupt important genome compositional features can have substantial phenotypic effects (95–97). For example, of 48 cefotaxime resistance mutations in TEM-1 β-lactamase that were identified from a randomly mutagenized pool of variants, 10 were silent (98). One mutation increased nitrocefin hydrolysis by 7.5-fold, probably by increasing the amount of functional TEM-1 (99). Our work on ΦX174 also shows that synonymous mutations can have massive fitness effects, as the virus can be completely attenuated by recoding with non-preferred, synonymous codons. Although we introduced many synonymous mutations in each recoded ΦX174 strain, the largest observed fitness impact of any single deoptimized fragment contained only 29 synonymous codon changes. These 29 synonymous changes resulted in a 50% decrease in fitness, which is a decrease of about 10 doublings per hour or about a 1000-fold change in the number of offspring. Aside from using codons that ensure the use of preferred tRNAs, organisms must also balance codon usage with codon pair biases, di-nucleotide preferences, mRNA structural motifs, restriction enzyme site avoidances, etc. Selection can act on many types of genome compositional features. In recoding ΦX174 with better or worse codons, we observed correlative changes in other compositional features (fig 6, supplementary table S4).

A good predictor of translational throughput is mRNA secondary structure (21). Tightly folded mRNAs are traversed by the ribosome more slowly than unfolded mRNAs, although faster translation is not necessarily beneficial. mRNA structure is used to slow translation where pausing is needed, most notably before protein structures that require co-translational folding (32, 33) and at the 5’ end of a transcript, where proper loading of a mRNA onto a ribosome foreshadows correct and efficient translation (29). Our data support the importance of mRNA folding stability for organismal fitness, as the stability of recoded ΦX174 genes is correlated with ΦX174 fitness (fig. 6b). Interestingly, the optimized ΦX174 strains almost always have increased mRNA folding stabilities and decreased fitness values, suggesting the folding stability of each mRNA may be near optimal in the wild-type virus. While it is unclear to us why the optimized strains have increased mRNA folding stabilities, departure from wild-type—either more or less stable—results in fitness declines. This suggests that wild-type ΦX174 sits at a peak in its fitness landscape.

### Combining mutations and epistasis

Genomes accumulate deleterious mutations over time. The detrimental effect of accumulating deleterious mutations is prevented by sex, recombination, and purifying selection which purge them from populations. In contrast to beneficial mutations which generally combine with diminishing returns (100, 101), the way that the individual effects of deleterious mutations combine is less well understood. Among many issues preventing these predictions is a paucity of empirical phenotype data for networks of deleterious mutations (102–104). This is especially true for combinations beyond two. A number of studies have investigated epistasis among pairs or triple sets of deleterious mutations, but the findings are mixed (105–107). Sometimes the combined effect is the sum of the individual effects (additive/no epistasis), sometimes it is less than predicted from the individual effects (antagonistic/positive epistasis) (108, 109), and sometimes it is more than predicted (synergistic/negative epistasis) (110). Among these three scenarios, antagonistic epistasis seems to be most common (104, 111). If one considers sign epistasis to be an extreme form of antagonistic epistasis, then more support is garnished for this model as a number of studies on deleterious mutations uncover some degree of deleterious mutations becoming beneficial in combination (112). Our data is novel in that it builds several complete combinatorial networks of deleterious mutations, but it does lack large samples sizes. Of the networks we built, only the one for gene A had a sufficient number of data points to reject poorly fitting models. For gene A, strong antagonistic epistasis was observed. Johnson *et al.* recently found that this type of epistasis is common among loss-of-function mutations in yeast. They called it “increasing cost epistasis” because a given deleterious mutation tends to have a greater cost on more fit backgrounds (113). For genes F and H, no models can be rejected, but the data suggest that mutations in gene F are additive while mutations in gene H combine synergistically. For the purposes of building synonymously recoded viruses for vaccines, it is promising to see gene A displaying antagonistic epistasis. With this type of epistasis where fitness flattens out, less trial and error should be required to build attenuated, but still viable, viruses.

### Synonymous virus genome recoding for vaccines

Synonymously recoding viral genomes has a potentially useful application in making live-attenuated vaccines. The antigenicity of synonymously attenuated viruses is maintained because the viral protein sequences remain unchanged. However, the process of choosing how many codons to change and what type of synonymous changes to implement is currently done without guiding principles. If fact, which synonymous features most strongly affect recoded viruses is debated (79, 114, 115, 115). Of the dozens of viruses that have been deoptimized, a minority of them measure compositional features different from the one being directly targeted for deoptimization. At the very least, we suggest that researchers must measure a variety of compositional features when designing deoptimized constructs. A better approach would be to develop construct design software that supports researchers to engineer deoptimized viral genes (see (116) for an example using codon shuffling). This software exists for optimizing genes for expression in host cells (117, 118) and may be co-opted for deoptimization purposes. In our experiments we made no effort to isolate changes to one type of compositional feature. In exploring this possibility, we found it difficult to generate sufficient deoptimization of one feature (CAI) while keeping other features (mfold, CpB, Shine-Dalgarno frequency) unchanged. Recently, Paff et al. demonstrated that promoter ablation attenuated T7 bacteriophage in a predictable manner (119). Combining these edits with previous codon deoptimized strains showed increased attenuation. Targeting intragenic attenuating mutations is a promising way to test how deleterious effects combine without the added complication of trying to isolate correlative compositional features.

Like many other studies, our data showed virus codon deoptimization is an effective way to generate attenuated viruses. In cell culture and animal studies, deoptimized viruses were shown to protect from viral challenge and were stable over small numbers of passages (60). However, much concern remains about the potential for attenuated viruses to recover virulence. It is therefore important to understand how many and what types of synonymous mutations can be made to viral genomes without completely ablating their ability to replicate in host cells. What viral genes should be attenuated? How many attenuating mutations should be made to the genome? What synonymous features should be targeted for deoptimization? In most studies to date, a limited number of deoptimized constructs (usually structural proteins) were tested. We showed that fitness decreases can be obtained by deoptimizing many of the ΦX174 genes, indicating that nonstructural genes may also be good targets for attenuation. One approach to avoid evolutionary reversion might be recoding multiple genes or entire viral genomes, balancing optimization and deoptimization to maintain sufficient virulence while increasing the genetic distance to wild type. This strategy could prevent recovery by mutation or by recombination with wild-type viruses. However, our work suggests that the effects of recoding will not be uniform across a genome. We found that the attenuating effects of recoding and the nature of epistatic interactions from combining fragments differ dramatically between genes.

## Acknowledgments

This study was supported by the National Institutes of Health grants P20 GM104420 and GM076040. We thank the Institute for Bioinformatic and Evolutionary Studies (IBEST) Genomics Resources Core for assistance with DNA sequencing. Sequences for all recoded ΦX174 genes are available on Genbank under accession numbers MN045299-MN045346.

## Methods and Methods

### Bacterial cultures and phage stocks

A laboratory strain of bacteriophage ΦX174 (GenBank accession number AF176034) was used in this study. All experiments were carried out using *E. coli* C as a host in modified Luria-Bertani media (10 g/l tryptone, 5 g/l Bacto yeast extract, 10 g/l NaCl, 2 mM CaCl2).

### Synthetic ΦX174 genomes

A phage assembly platform for ΦX174 was devised following (120). The ΦX174 chromosome was divided into 15 genomic fragments designed to avoid host cell toxicity by separating genes from their promoters and breaking large genes into multiple segments. Each segment is flanked by unique five nucleotide overlaps of WT ΦX174 sequence so that they can be amplified from the ancestral ΦX174 using PCR primers designed to incorporate terminal BsmB1 restriction sites. Amplicons were cloned into pCR2.1 using the Invitrogen TOPO TA cloning system (Life Technologies, Grand Island, NY). We pooled plasmid DNA containing all 15 of the phage DNA fragments in equimolar amounts and digested them with BsmB1 (Fermentas Fast Digest, Life Technologies, Grand Island, NY) for 30 minutes to 1 hour at 37°C. The digested plasmids were subjected to agarose gel electrophoresis for 10 to 15 minutes using a 1.2% agarose gel to separate the vector from the inserts. The inserts were excised from the gel, purified using the GeneJET gel extraction kit (Fermentas), ligated overnight at 14°C with T4 DNA ligase (Promega Corporation, Madison, WI), and transformed by electroporation into 100 μl competent *E. coli* C cells. The transformation mix was resuspended with 1 ml of ΦLB and either plated immediately or incubated for about 20 minutes at 37°C to allow for one viral burst. The ΦLB was added to 3 ml of ΦLB top agar and plated onto a ΦLB agar plate. After four to five hours of incubation at 37°C, recombinant phage plaques were visible and plates were removed from the incubator. To verify that the recombinant phage encoded the intended sequence, the resulting phage genome was sequenced in its entirety as previously described (121).

### Codon deoptimization of ΦX174

Codon deoptimized and optimized fragments were synthesized by in-house at the University of Texas at Austin, Applied Research Laboratories’ Gene Synthesis Facility or purchased from Biomatik USA, LLC (Wilmington, DE) according to the codon usage of five representative *E. coli* genomes (*E. coli* 536, *E. coli* UT 189, *E. coli* O157:H7 str. Sakai, *E. coli* O157:H7EDL933, and *E. coli* CFT073). Codon usage was calculated by averaging each codon’s usage frequency in CDS of these *E. coli* genomes. Each wild-type ΦX174 codon that could be changed to a more or less frequently used codon was changed. Some coding regions were not modified; these were either identified a priori or excluded based on failure of a genome construct to yield viable phage. Unmodified regions in our final dataset include overlapping reading frames, promoter regions that occurred within other reading frames, and the region from A4299-A4328 which encodes the ΦX174 origin of replication. In addition, approximately 50 bases surrounding the initiation codon (AUG) of each gene were left unmodified to assure efficient translation initiation. Unsuccessful attempts to create live virus using synthesized fragments were repeated at least three times, then passaged in liquid culture for 24 hours to allow for recovery mutants to arise.

### Measuring viral fitness

Fitness assays and fitness calculations were performed as previously described (27). The assay is a determination of growth rate at low MOI in 10 mL LB and is carried out at 37°C. Host cells were prepared by growing to ~10^8^ cells/mL and aliquoted into 8.5 mL of warm LB just prior to adding phage. Phage fitness is expressed as the log2 fold increase in the total number of phage per hour. All measurements were done in triplicate. At 45min, virus titers were done on LB-agar plates with 0.3% top agar.

### Calculating genome statistics

Calculations of genome statistics were done as follows: Codon Adaptation Index (CAI) was calculated using the seqinr cai() function in R which uses the Sharp and Li 1987 method and the *E. coli* codon usage table. An alternative form of CAI was also calculated using an updated method outlined in Xia 2007. The number of effective codons (Nc) was calculated following (85). The Index of translational elongation (I_TE_) was calculated according to (83). The Starvation codon adaptation index (sCAI) was calculated according to (86). This measure scores genes by how susceptible their codons are to a scarcity of amino-acylated tRNAs. The number of Shine-Dalgarno motifs with binding values > 0 were counted in any given sequence and included in the model. The strength of the Shine-Dalgarno sequences was also considered by including a per-codon average binding strength of all Shine-Dalgarno motifs in a gene (sum of binding strengths/gene gene length).

For Figure 3, all protein coding sequences were parsed from the *E. coli* O103:H2 genome (GCA_000010745.1) and CAI was calculated as described above. The list of the most highly expressed genes is from (122).

### Analysis of Epistasis

We analyzed the network fitness data from genes A, F, and H using the Stickbreaker R package (88) and functions therein. This package fits such data to the additive, multiplicative, and stickbreaking models. While the additive and multiplicative models assume a mutation (a recoded block in this context) changes background fitness by a difference or a factor (respectively), the stickbreaking model assumes a mutation’s effect is scaled by the distance between the background and a fitness boundary. For fitting the stickbreaking model, we could not obtain reasonable estimates for the fitness boundary from the data (beneficial mutations are much more useful for estimating the boundary than deleterious ones). Instead, we assumed a fitness boundary of 24.5 dbl/hr (wildtype has fitness of 20.5); using a larger fitness boundary simply makes the sticking model more and more like the additive model. Relative fit (posterior probabilities) was calculated following the methods in (88). To estimate the absolute goodness of fit, we used parametric bootstrap. Specifically, for each gene and each model, we extracted the observed effect of each block on each background it appeared on. For each recoded fragment, we then regressed the background’s fitness against the fitness effect by fitting a simple linear model and obtained a p-value associated with a slope of zero (illustrated for gene A in fig 5). When a model is correct, the slope of this line is expected to be zero. For a given gene and model, we take the sum of the logs of the p-values, P_obs_, as a summary statistic. Noticed that the data points involved in these regressions were not independent and, as such, the p-values were not valid. We accounted for this by simulating 10,000 datasets (using the estimated coefficient of each block and the estimated Gaussian noise parameter, that captures both experimental noise and variation from model expectations). For each simulated dataset, we repeated the regression for each block and combined across blocks to obtain a summary P_sim_. Across 10,000 simulations, this generated the approximate distribution of P when the model is correct. We then located P_obs_ in this distribution and calculated the p-value as the proportion of simulations where P_sim_ < P_obs_.

