## supplementary materials for "ΦX174 Attenuation by Whole Genome Codon Deoptimization"

**Supplementary fig. S1 -** Using parametric bootstrap, the distribution of the sum of the recoded fragment's individual p-values under the additive model (fig. 5b) was simulated. The p-value is estimated by the proportion of simulations where the sum of logs is ≤ the observed value.

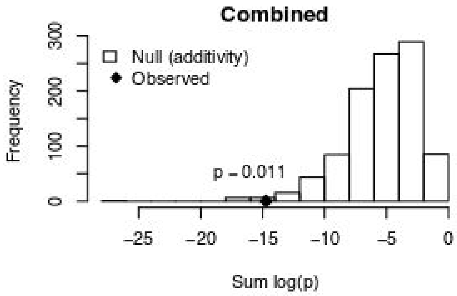


**Supplementary fig. S2 -** Regression of each recoded fragment's fitness effect under the multiplicative model against background fitness. The p-values of each regression are combined by taking the sum of their logs. Using parametric bootstrap, the distribution of this sum was simulated (bottom panel). The p-value is estimated by the proportion of simulations where the sum of logs is ≤ the observed value.

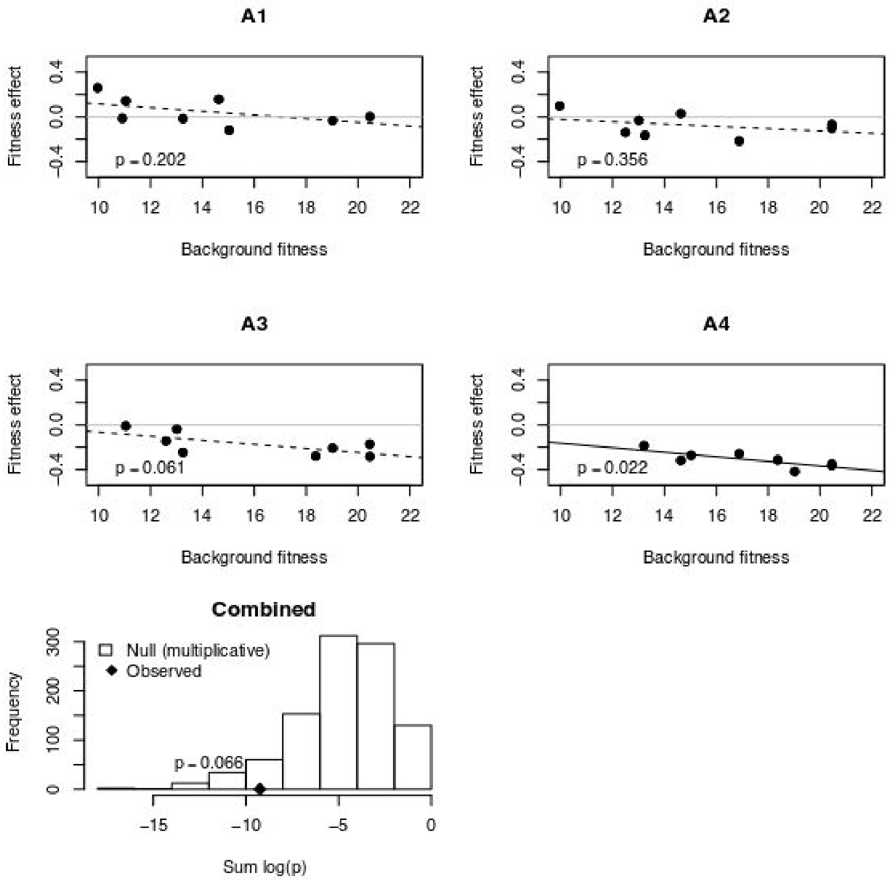

**Supplementary fig. S3 -** Regression of each recoded fragment's fitness effect under the stickbreaking (synergistic) model against background fitness. The p-values of each regression are combined by taking the sum of their logs. Using parametric bootstrap, the distribution of this sum was simulated (bottom panel). The p-value is estimated by the proportion of simulations where the sum of logs is ≤ the observed value.

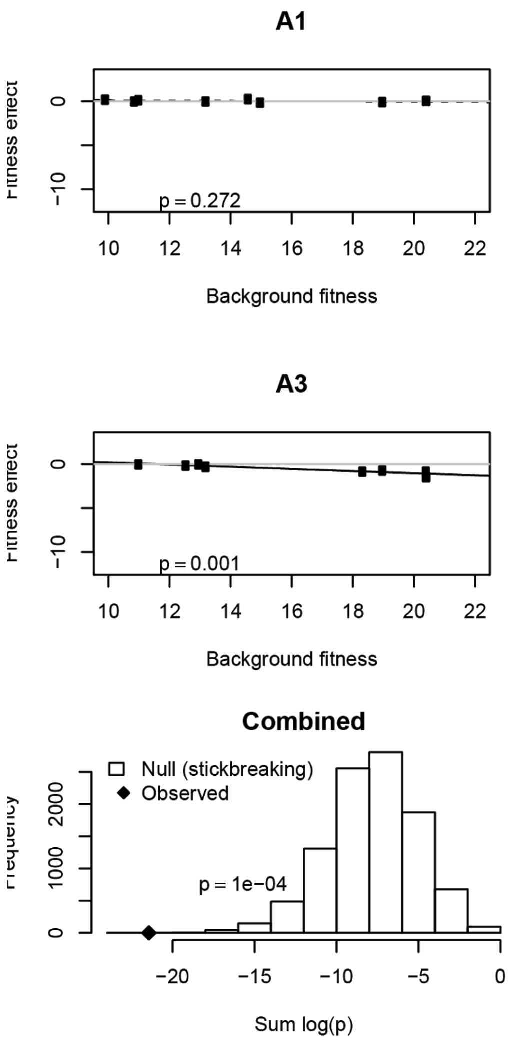

**Supplementary table 1.** Codon preference statistics for wildtype and codon edited genes. iSD is the per codon average Shine-Dalgarno motif binding strength across a gene.

| **Genotype** | **GC** | **RSCU** | **CAI (Sharp)** | **CAI (Xia 2007)** | **$I_{TE}$** | **sCAI** | **iSD** | **tAI** | **tAI (normalized)** | **CPB** | **mfold** |
|---|---|---|---|---|---|---|---|---|---|---|---|
| A1 | 0.46 | 0.679 | 0.270 | 0.456 | 0.689 | 9.6 | -0.267 | 0.192 | 0.487 | 2.60 | -457.6 |
| A1_A2 | 0.46 | 0.418 | 0.174 | 0.328 | 0.666 | 10.6 | -0.328 | 0.192 | 0.487 | -1.12 | -441.6 |
| A1_A2_A3 | 0.46 | 0.289 | 0.134 | 0.285 | 0.663 | 10.8 | -0.376 | 0.198 | 0.504 | -3.45 | -429.2 |
| A1_A2_A3_A4 | 0.46 | 0.244 | 0.114 | 0.251 | 0.649 | 10.6 | -0.382 | 0.200 | 0.507 | -4.94 | -428.9 |
| A1_A2_A4 | 0.46 | 0.354 | 0.149 | 0.289 | 0.652 | 10.4 | -0.334 | 0.193 | 0.490 | -2.61 | -436.3 |
| A1_A3 | 0.46 | 0.469 | 0.206 | 0.396 | 0.686 | 9.8 | -0.315 | 0.199 | 0.504 | 0.26 | -442.1 |
| A1_A3_A4 | 0.46 | 0.396 | 0.176 | 0.349 | 0.671 | 9.6 | -0.321 | 0.200 | 0.508 | -1.23 | -435.8 |
| A1_A4 | 0.46 | 0.574 | 0.230 | 0.402 | 0.674 | 9.4 | -0.273 | 0.193 | 0.490 | 1.11 | -450.5 |
| A1_opt | 0.45 | 0.695 | 0.272 | 0.486 | 0.710 | 9.6 | -0.234 | 0.205 | 0.506 | 2.89 | -472.9 |
| A2 | 0.46 | 0.472 | 0.195 | 0.358 | 0.667 | 10.7 | -0.316 | 0.191 | 0.486 | -0.29 | -438.9 |
| A2_A3 | 0.46 | 0.326 | 0.150 | 0.311 | 0.664 | 10.9 | -0.364 | 0.198 | 0.503 | -2.63 | -426.4 |
| A2_A3_A4 | 0.46 | 0.276 | 0.128 | 0.274 | 0.650 | 10.7 | -0.370 | 0.199 | 0.506 | -4.12 | -425.8 |
| A2_A4 | 0.46 | 0.399 | 0.167 | 0.315 | 0.653 | 10.5 | -0.322 | 0.193 | 0.489 | -1.78 | -433.6 |
| A2_opt | 0.46 | 0.761 | 0.306 | 0.529 | 0.709 | 10.0 | -0.231 | 0.218 | 0.538 | 5.51 | -469.9 |
| A3 | 0.45 | 0.529 | 0.231 | 0.431 | 0.687 | 9.9 | -0.303 | 0.198 | 0.503 | 1.09 | -439.9 |
| A3_A4 | 0.46 | 0.447 | 0.197 | 0.380 | 0.672 | 9.7 | -0.309 | 0.200 | 0.507 | -0.40 | -433.7 |
| A3_opt | 0.47 | 0.781 | 0.318 | 0.535 | 0.715 | 10.4 | -0.229 | 0.216 | 0.533 | 3.93 | -486.6 |
| A4 | 0.45 | 0.648 | 0.258 | 0.438 | 0.675 | 9.5 | -0.261 | 0.193 | 0.489 | 1.93 | -446.6 |
| Awt | 0.45 | 0.767 | 0.302 | 0.497 | 0.690 | 9.7 | -0.255 | 0.192 | 0.486 | 3.42 | -455.4 |
| C | 0.44 | 0.373 | 0.164 | 0.342 | 0.654 | 13.7 | -0.280 | 0.200 | 0.508 | -1.58 | -65.3 |
| C_opt | 0.47 | 0.704 | 0.301 | 0.523 | 0.645 | 12.0 | -0.167 | 0.233 | 0.596 | 5.35 | -67.9 |
| Cwt | 0.42 | 0.486 | 0.183 | 0.414 | 0.649 | 13.4 | -0.244 | 0.181 | 0.461 | -0.07 | -69.1 |
| F1 | 0.45 | 0.633 | 0.254 | 0.424 | 0.695 | 11.3 | -0.229 | 0.190 | 0.513 | 2.75 | -370.9 |
| F1_F2 | 0.45 | 0.360 | 0.153 | 0.289 | 0.649 | 11.1 | -0.273 | 0.188 | 0.507 | -0.87 | -349.4 |
| F1_F2_F3 | 0.45 | 0.180 | 0.082 | 0.170 | 0.593 | 10.9 | -0.332 | 0.191 | 0.516 | -5.82 | -302.3 |
| F1_F3 | 0.45 | 0.318 | 0.136 | 0.251 | 0.639 | 11.2 | -0.288 | 0.193 | 0.522 | -2.12 | -338.1 |
| F1_opt | 0.47 | 0.993 | 0.389 | 0.601 | 0.700 | 11.9 | -0.199 | 0.200 | 0.539 | 6.62 | -404.6 |
| F2 | 0.45 | 0.547 | 0.222 | 0.380 | 0.672 | 12.4 | -0.237 | 0.188 | 0.508 | 1.92 | -357.7 |
| F2_F3 | 0.46 | 0.275 | 0.119 | 0.225 | 0.618 | 12.3 | -0.295 | 0.191 | 0.517 | -2.95 | -315.2 |
| F2_opt | 0.49 | 1.074 | 0.436 | 0.644 | 0.711 | 13.2 | -0.197 | 0.209 | 0.564 | 7.82 | -407.4 |
| F3 | 0.45 | 0.483 | 0.198 | 0.331 | 0.662 | 12.5 | -0.251 | 0.193 | 0.523 | 0.67 | -346.2 |
| F3_opt | 0.49 | 1.066 | 0.431 | 0.634 | 0.694 | 13.1 | -0.213 | 0.211 | 0.571 | 8.10 | -422.6 |
| Fwt | 0.45 | 0.961 | 0.370 | 0.558 | 0.720 | 12.7 | -0.192 | 0.190 | 0.514 | 5.54 | -374.2 |

| | | | | | | | | | | |
|---|---|---|---|---|---|---|---|---|---|---|
| G1 | 0.43 | 0.375 | 0.162 | 0.251 | 0.629 | 9.8 | -0.202 | 0.191 | 0.521 | -0.96 | -118.1 |
| G1_opt | 0.53 | 1.107 | 0.486 | 0.743 | 0.688 | 12.1 | -0.256 | 0.211 | 0.575 | 8.58 | -187.2 |
| G2 | 0.41 | 0.536 | 0.211 | 0.364 | 0.630 | 10.8 | -0.211 | 0.161 | 0.438 | 2.05 | -130.9 |
| G2_opt | 0.44 | 0.815 | 0.319 | 0.529 | 0.648 | 10.6 | -0.215 | 0.178 | 0.483 | 5.12 | -141.3 |
| Gwt | 0.42 | 0.788 | 0.302 | 0.504 | 0.660 | 10.5 | -0.194 | 0.167 | 0.455 | 4.84 | -133.2 |
| H1 | 0.45 | 0.523 | 0.217 | 0.338 | 0.652 | 10.4 | -0.379 | 0.194 | 0.464 | 1.87 | -289.4 |
| H1_H2 | 0.44 | 0.338 | 0.142 | 0.243 | 0.631 | 10.3 | -0.413 | 0.188 | 0.450 | -2.70 | -273.9 |
| H1_H2_H3 | 0.44 | 0.257 | 0.109 | 0.192 | 0.646 | 10.3 | -0.447 | 0.191 | 0.457 | -5.55 | -259.8 |
| H1_H3 | 0.45 | 0.398 | 0.166 | 0.267 | 0.667 | 10.4 | -0.413 | 0.197 | 0.471 | -1.01 | -282.3 |
| H1_opt | 0.51 | 0.974 | 0.425 | 0.638 | 0.699 | 11.0 | -0.331 | 0.229 | 0.548 | 8.69 | -338.3 |
| H2 | 0.44 | 0.550 | 0.227 | 0.385 | 0.662 | 11.9 | -0.369 | 0.194 | 0.465 | 0.98 | -305.5 |
| H2_H3 | 0.44 | 0.411 | 0.172 | 0.306 | 0.677 | 11.9 | -0.403 | 0.198 | 0.473 | -1.88 | -298.7 |
| H2_opt | 0.49 | 1.003 | 0.438 | 0.677 | 0.722 | 10.2 | -0.315 | 0.226 | 0.540 | 5.86 | -334.2 |
| H3 | 0.45 | 0.647 | 0.265 | 0.422 | 0.699 | 12.1 | -0.369 | 0.204 | 0.488 | 2.67 | -314.5 |
| H3_opt | 0.48 | 0.886 | 0.368 | 0.570 | 0.681 | 12.2 | -0.348 | 0.212 | 0.507 | 6.73 | -337.7 |
| Hwt | 0.45 | 0.851 | 0.346 | 0.534 | 0.684 | 12.1 | -0.335 | 0.201 | 0.480 | 5.55 | -305.8 |
| J | 0.50 | 0.126 | 0.058 | 0.195 | 0.629 | 9.5 | -0.730 | 0.201 | 0.468 | -2.84 | -25.9 |
| J_opt | 0.64 | 1.551 | 0.674 | 0.841 | 0.762 | 13.5 | -0.324 | 0.338 | 0.785 | 12.62 | -54.8 |
| Jwt | 0.50 | 1.172 | 0.483 | 0.622 | 0.838 | 14.1 | -0.397 | 0.231 | 0.537 | 8.73 | -34.8 |

**Supplementary table 2 -** Edits are evenly distributed across recoded ΦX174 genes. The length of recoded genome segments, number of altered nucleotides, and number of altered codons are shown. The proportion of nucleotides edited (relative to the length of the gene or the length of the recoded fragment) are in parentheses.

| | | Deoptimized | | Optimized | |
|---|---|---|---|---|---|
| **Fragment** | **Length (bp)** | **# bp changed** | **# codons changed** | **# bp changed** | **# codons changed** |
| A1 | 90 (0.06) | 19 (0.21) | 18 | 27 (0.30) | 21 |
| A2 | 489 (0.32) | 99 (0.20) | 88 | 84 (0.17) | 75 |
| A3 | 306 (0.20) | 80 (0.26) | 64 | 69 (0.23) | 65 |
| A4 | 153 (0.10) | 37 (0.24) | 32 | 38 (0.25) | 32 |
| C | 144 (0.55) | 21 (0.15) | 20 | 40 (0.28) | 31 |
| J | 96 (0.82) | 30 (0.32) | 24 | 22 (0.23) | 20 |
| F1 | 267 (0.21) | 72 (0.27) | 65 | 68 (0.26) | 60 |
| F2 | 441 (0.34) | 115 (0.26) | 103 | 102 (0.23) | 87 |
| F3 | 552 (0.43) | 165 (0.30) | 146 | 128 (0.23) | 100 |
| G1 | 390 (0.74) | 108 (0.28) | 100 | 112 (0.29) | 86 |
| G2 | 117 (0.22) | 31 (0.27) | 29 | 21 (0.18) | 17 |
| H1 | 336 (0.34) | 89 (0.27) | 83 | 92 (0.27) | 75 |
| H2 | 393 (0.40) | 76 (0.19) | 72 | 98 (0.25) | 85 |
| H3 | 240 (0.24) | 61 (0.26) | 57 | 61 (0.26) | 47 |

**Supplementary table 3 -** Fitness values for all constructs by date and plaque number.

| Genotype | Date | Plaque | Fitness | Model Fitness |
|---|---|---|---|---|
| A1 | 12/28/2016 | 2 | 23.5 | 21.7 |
| A1 | 1/5/2017 | 1 | 22.3 | 21.7 |
| A1 | 1/10/2017 | 3 | 23.2 | 21.7 |
| A1_A2 | 12/20/2016 | 1 | 19.9 | 19.6 |
| A1_A2 | 12/28/2016 | 2 | 20.8 | 19.6 |
| A1_A2 | 1/3/2017 | 3 | 21.8 | 19.6 |
| A1_A2_A3 | 12/20/2016 | 1 | 14.3 | 14.4 |
| A1_A2_A3 | 12/28/2016 | 2 | 18.1 | 14.4 |
| A1_A2_A3 | 1/4/2017 | 3 | 15.6 | 14.4 |
| A1_A2_A3 | 1/18/2018 | 3 | 14.8 | 14.4 |
| A1_A2_A3 | 1/25/2018 | 3 | 13.0 | 14.4 |
| A1_A2_A3 | 1/25/2018 | 1 | 16.5 | 14.4 |
| A1_A2_A3 | 12/28/2017 | 1 | 15.6 | 14.4 |
| A1_A2_A3 | 1/12/2018 | 1 | 14.9 | 14.4 |
| A1_A2_A3 | 1/18/2018 | 1 | 17.9 | 14.4 |
| A1_A2_A3 | 1/25/2018 | 2 | 13.0 | 14.4 |
| A1_A2_A3 | 12/28/2017 | 2 | 16.0 | 14.4 |
| A1_A2_A3 | 1/12/2018 | 2 | 14.4 | 14.4 |
| A1_A2_A3 | 1/18/2018 | 2 | 17.4 | 14.4 |
| A1_A2_A3 | 12/28/2017 | 3 | 15.8 | 14.4 |
| A1_A2_A3 | 1/12/2018 | 3 | 17.0 | 14.4 |
| A1_A2_A3_A4 | 1/4/2017 | 3 | 13.9 | 11.5 |
| A1_A2_A3_A4 | 12/30/2016 | 2 | 12.2 | 11.5 |
| A1_A2_A3_A4 | 12/20/2016 | 1 | 10.6 | 11.5 |
| A1_A2_A4 | 1/10/2017 | 2 | 13.7 | 13.6 |
| A1_A2_A4 | 12/20/2016 | 1 | 14.2 | 13.6 |
| A1_A2_A4 | 1/4/2017 | 3 | 16.1 | 13.6 |
| A1_A3 | 1/3/2017 | 3 | 22.5 | 17.9 |
| A1_A3 | 1/12/2017 | 1 | 17.8 | 17.9 |
| A1_A3 | 1/12/2017 | 2 | 16.2 | 17.9 |
| A1_A3_A4 | 12/30/2016 | 2 | 13.4 | 13.4 |
| A1_A3_A4 | 1/4/2017 | 3 | 13.8 | 13.4 |
| A1_A3_A4 | 1/5/2017 | 1 | 15.6 | 13.4 |
| A1_A4 | 12/20/2016 | 1 | 14.8 | 14.2 |
| A1_A4 | 12/28/2016 | 2 | 15.2 | 14.2 |
| A1_A4 | 1/4/2017 | 3 | 15.9 | 14.2 |
| A1_opt | 7/30/2012 | 1 | 19.4 | 19.8 |
| A1_opt | 7/30/2012 | 3 | 18.8 | 19.8 |

| | | | | |
|---|---|---|---|---|
| A1_opt | 4/6/2012 | 4 | 20.4 | 19.8 |
| A1_opt | 7/26/2012 | 2 | 17.1 | 19.8 |
| A2 | 12/19/2016 | 1 | 22.1 | 20.5 |
| A2 | 12/28/2016 | 2 | 21.4 | 20.5 |
| A2 | 1/3/2017 | 3 | 22.6 | 20.5 |
| A2_A3 | 1/4/2017 | 3 | 16.9 | 15.9 |
| A2_A3 | 12/20/2016 | 1 | 16.3 | 15.9 |
| A2_A3 | 1/12/2017 | 2 | 16.9 | 15.9 |
| A2_A3_A4 | 12/20/2016 | 1 | 10.6 | 11.6 |
| A2_A3_A4 | 12/30/2016 | 2 | 11.4 | 11.6 |
| A2_A3_A4 | 1/4/2017 | 3 | 15.1 | 11.6 |
| A2_A4 | 1/10/2017 | 2 | 13.6 | 12.1 |
| A2_A4 | 12/20/2016 | 1 | 11.7 | 12.1 |
| A2_A4 | 1/4/2017 | 3 | 14.0 | 12.1 |
| A2_opt | 3/1/2012 | 3 | 18.0 | 19.4 |
| A2_opt | 3/1/2012 | 4 | 19.5 | 19.4 |
| A2_opt | 3/1/2012 | 1 | 20.0 | 19.4 |
| A2_opt | 3/1/2012 | 2 | 19.5 | 19.4 |
| A3 | 12/28/2016 | 2 | 18.4 | 15.9 |
| A3 | 1/5/2017 | 1 | 16.4 | 15.9 |
| A3 | 1/10/2017 | 3 | 16.6 | 15.9 |
| A3_A4 | 1/4/2017 | 3 | 11.9 | 11.0 |
| A3_A4 | 1/10/2017 | 2 | 13.7 | 11.0 |
| A3_A4 | 12/20/2016 | 1 | 10.5 | 11.0 |
| A3_opt | 7/26/2012 | 1 | 10.4 | 13.7 |
| A3_opt | 7/30/2012 | 2 | 13.9 | 13.7 |
| A3_opt | 10/5/2012 | 6 | 15.2 | 13.7 |
| A3_opt | 11/21/2012 | 8 | 13.3 | 13.7 |
| A3_opt | 8/1/2012 | 5 | 12.3 | 13.7 |
| A3_opt | 11/20/2012 | 7 | 12.2 | 13.7 |
| A3_opt | 8/1/2012 | 4 | 11.7 | 13.7 |
| A3_opt | 7/30/2012 | 3 | 13.9 | 13.7 |
| A4 | 1/3/2017 | 3 | 17.2 | 14.5 |
| A4 | 12/20/2016 | 1 | 13.6 | 14.5 |
| A4 | 12/28/2016 | 2 | 16.3 | 14.5 |
| F1 | 12/22/2016 | 2 | 20.0 | 18.7 |
| F1 | 12/30/2016 | 3 | 20.3 | 18.7 |
| F1 | 1/12/2017 | 1 | 18.2 | 18.7 |
| F1_F2 | 12/19/2016 | 1 | 20.0 | 17.7 |
| F1_F2 | 12/30/2016 | 3 | 16.7 | 17.7 |
| F1_F2 | 1/12/2017 | 2 | 18.8 | 17.7 |

| | | | | |
|---|---|---|---|---|
| F1_F2_F3 | 1/10/2017 | 1 | 9.1 | 8.0 |
| F1_F2_F3 | 1/3/2017 | 3 | 11.1 | 8.0 |
| F1_F2_F3 | 1/5/2017 | 2 | 7.6 | 8.0 |
| F1_F3 | 1/5/2017 | 2 | 11.7 | 11.1 |
| F1_F3 | 12/19/2016 | 1 | 12.7 | 11.1 |
| F1_F3 | 1/10/2017 | 3 | 12.2 | 11.1 |
| F1_opt | 3/1/2012 | 3 | 20.0 | 20.2 |
| F1_opt | 3/1/2012 | 2 | 21.0 | 20.2 |
| F1_opt | 3/1/2012 | 1 | 20.0 | 20.2 |
| F1_opt | 3/1/2012 | 4 | 19.8 | 20.2 |
| F2 | 12/30/2016 | 3 | 19.8 | 19.7 |
| F2 | 12/19/2016 | 1 | 20.5 | 19.7 |
| F2 | 12/22/2016 | 2 | 21.6 | 19.7 |
| F2_F3 | 12/19/2016 | 1 | 12.0 | 10.2 |
| F2_F3 | 12/22/2016 | 2 | 11.0 | 10.2 |
| F2_F3 | 1/10/2017 | 3 | 11.3 | 10.2 |
| F2_opt | 3/1/2012 | 4 | 21.0 | 20.7 |
| F2_opt | 3/1/2012 | 1 | 19.4 | 20.7 |
| F2_opt | 3/1/2012 | 2 | 21.5 | 20.7 |
| F2_opt | 3/1/2012 | 3 | 21.0 | 20.7 |
| F3 | 1/5/2017 | 1 | 14.2 | 14.3 |
| F3 | 12/22/2016 | 2 | 15.4 | 14.3 |
| F3 | 1/12/2017 | 3 | 16.0 | 14.3 |
| F3_opt | 11/4/2011 | 5 | 20.0 | 19.2 |
| F3_opt | 11/1/2011 | 3 | 17.8 | 19.2 |
| F3_opt | 11/4/2011 | 6 | 19.4 | 19.2 |
| F3_opt | 10/31/2011 | 2 | 18.8 | 19.2 |
| F3_opt | 11/1/2011 | 4 | 17.3 | 19.2 |
| F3_opt | 10/31/2011 | 1 | 18.7 | 19.2 |
| F3_opt | 11/7/2011 | 7 | 18.8 | 19.2 |
| F3_opt | 11/10/2011 | 9 | 18.9 | 19.2 |
| F3_opt | 11/10/2011 | 10 | 19.5 | 19.2 |
| F3_opt | 11/7/2011 | 8 | 19.3 | 19.2 |
| G1 | 12/22/2016 | 2 | 11.2 | 10.4 |
| G1 | 1/3/2017 | 3 | 12.0 | 10.4 |
| G1 | 1/5/2017 | 1 | 11.8 | 10.4 |
| G1_opt | 3/22/2012 | 3 | 15.4 | 18.6 |
| G1_opt | 3/26/2012 | 2 | 18.7 | 18.6 |
| G1_opt | 3/19/2012 | 4 | 17.9 | 18.6 |
| G1_opt | 3/28/2012 | 1 | 17.0 | 18.6 |
| G2 | 1/3/2017 | 3 | 10.2 | 9.3 |

| | | | | |
|---|---|---|---|---|
| G2 | 12/22/2016 | 2 | 11.6 | 9.3 |
| G2 | 1/12/2017 | 1 | 9.6 | 9.3 |
| H1 | 5/21/2017 | 1 | 24.3 | 21.9 |
| H1 | 5/21/2017 | 2 | 26.4 | 21.9 |
| H1 | 6/9/2017 | 1 | 26.3 | 21.9 |
| H1 | 12/19/2016 | 1 | 21.8 | 21.9 |
| H1 | 6/9/2017 | 2 | 24.4 | 21.9 |
| H1 | 5/21/2017 | 3 | 25.9 | 21.9 |
| H1 | 1/3/2017 | 3 | 21.8 | 21.9 |
| H1 | 6/9/2017 | 3 | 25.0 | 21.9 |
| H1 | 12/22/2016 | 2 | 22.5 | 21.9 |
| H1_H2 | 6/9/2017 | 3 | 16.1 | 13.7 |
| H1_H2 | 5/21/2017 | 1 | 16.6 | 13.7 |
| H1_H2 | 5/21/2017 | 3 | 18.3 | 13.7 |
| H1_H2 | 6/9/2017 | 1 | 17.0 | 13.7 |
| H1_H2 | 5/21/2017 | 2 | 16.2 | 13.7 |
| H1_H2 | 6/9/2017 | 2 | 15.8 | 13.7 |
| H1_H2_H3 | 5/21/2017 | 2 | 14.3 | 13.4 |
| H1_H2_H3 | 5/21/2017 | 1 | 16.1 | 13.4 |
| H1_H2_H3 | 6/9/2017 | 3 | 21.2 | 13.4 |
| H1_H2_H3 | 5/21/2017 | 3 | 16.4 | 13.4 |
| H1_H2_H3 | 6/9/2017 | 1 | 15.0 | 13.4 |
| H1_H2_H3 | 6/9/2017 | 2 | 14.7 | 13.4 |
| H1_H3 | 1/3/2017 | 3 | 18.5 | 18.9 |
| H1_H3 | 6/9/2017 | 1 | 21.7 | 18.9 |
| H1_H3 | 12/28/2016 | 2 | 19.2 | 18.9 |
| H1_H3 | 5/21/2017 | 3 | 23.3 | 18.9 |
| H1_H3 | 5/21/2017 | 2 | 23.1 | 18.9 |
| H1_H3 | 6/9/2017 | 3 | 22.6 | 18.9 |
| H1_H3 | 5/21/2017 | 1 | 23.2 | 18.9 |
| H1_H3 | 12/19/2016 | 1 | 18.1 | 18.9 |
| H1_H3 | 6/9/2017 | 2 | 22.6 | 18.9 |
| H2 | 1/10/2017 | 3 | 21.7 | 20.6 |
| H2 | 12/28/2016 | 2 | 22.4 | 20.6 |
| H2 | 12/19/2016 | 1 | 21.7 | 20.6 |
| H2_H3 | 1/3/2017 | 3 | 20.6 | 19.7 |
| H2_H3 | 12/19/2016 | 1 | 22.0 | 19.7 |
| H2_H3 | 1/12/2017 | 2 | 20.1 | 19.7 |
| H2_opt | 3/1/2012 | 4 | 20.2 | 20.4 |
| H2_opt | 3/1/2012 | 2 | 21.5 | 20.4 |
| H2_opt | 3/1/2012 | 1 | 20.0 | 20.4 |

| | | | | |
|---|---|---|---|---|
| H2_opt | 3/1/2012 | 3 | 19.8 | 20.4 |
| H3 | 1/10/2017 | 3 | 22.5 | 21.5 |
| H3 | 1/12/2017 | 1 | 22.5 | 21.5 |
| H3 | 1/12/2017 | 2 | 22.2 | 21.5 |
| H3_opt | 4/6/2012 | 4 | 20.9 | 19.7 |
| H3_opt | 7/30/2012 | 1 | 19.6 | 19.7 |
| H3_opt | 7/26/2012 | 2 | 15.4 | 19.7 |
| H3_opt | 7/30/2012 | 3 | 19.5 | 19.7 |

**Supplementary table 4 -** General linearized model showing correlation between fitness effect and various genomic features. See materials and methods for details on how the values were calculated. We included both the absolute values as well as the ratio between wildtype and the recoded variants.

| **metric** | **$R^2$** | **pvalue** | **AIC** | **dAIC** | **padjust** |
|---|---|---|---|---|---|
| RNA folding (mfold) | 0.67 | 6.6E-13 | 267.5 | 0.0 | 0.02 |
| FracEdited | 0.59 | 1.9E-10 | 279.8 | 12.3 | 0.38 |
| #ntsChanged | 0.59 | 1.9E-10 | 279.8 | 12.3 | 0.38 |
| CPB | 0.58 | 3.4E-10 | 282.1 | 14.5 | 0.44 |
| CaiXia2007Ratio | 0.58 | 4.8E-10 | 283.1 | 15.5 | 0.47 |
| IteIsoRatio | 0.57 | 9.5E-10 | 284.1 | 16.6 | 0.17 |
| CaiSharp1987Ratio | 0.57 | 6.9E-10 | 284.5 | 17.0 | 0.53 |
| IteIso | 0.56 | 1.2E-09 | 284.8 | 17.3 | 0.18 |
| #InframeShineDalgaros | 0.52 | 1.4E-08 | 289.6 | 22.1 | 0.50 |
| CaiXia2007 | 0.50 | 4.1E-08 | 293.1 | 25.6 | 0.58 |
| nSD | 0.48 | 1.3E-07 | 294.5 | 27.0 | 0.48 |
| CaiSharp1987 | 0.47 | 1.6E-07 | 295.7 | 28.2 | 0.63 |
| RSCU | 0.47 | 1.9E-07 | 296.0 | 28.4 | 0.63 |
| #ShineDalgaros | 0.44 | 7.9E-07 | 299.3 | 31.7 | 0.59 |
| tAI | 0.38 | 9.7E-06 | 302.2 | 34.6 | 0.07 |
| empirical | 0.16 | 2.5E-02 | 318.8 | 51.3 | 0.57 |
| CpbRatio | 0.02 | 8.2E-01 | 327.7 | 60.2 | 0.84 |
